# Multiple Toll-Spätzle Pathways in *Drosophila melanogaster* Immunity

**DOI:** 10.1101/420679

**Authors:** Munmun Chowdhury, Chun-Feng Li, Zhen He, Yuzhen Lu, Xusheng Liu, Yufeng Wang, Y. Tony Ip, Michael R. Strand, Xiao-Qiang Yu

## Abstract

The *Drosophila melanogaster* Toll-Spätzle pathway plays an important role in development and immunity. *Drosophila* genome encodes nine Toll receptors and six Spätzle (Spz) proteins, and only the canonical Toll-Spz (Toll-1-Spz-1) pathway has been well investigated. In this study, we compared the nine *Drosophila* Tolls and showed that similarly to Toll, Toll-7 also strongly activated *drosomycin* promoter. Importantly, we showed that both Toll and Toll-7 interacted with Spz, Spz-2 and Spz-5, and co-expression of Toll or Toll-7 with Spz, Spz-2 and Spz-5 activated the *drosomycin* promoter. Furthermore, Toll and Toll-7 both recognized *vesicular stomatitis* virus (VSV) by binding to the VSV glycoprotein. Septic infection in Toll and Toll-7 mutant flies suggested that Toll and Toll-7 differentially affected defense responses in adult males and females after systemic infection by *Enterococcus faecalis*, *Pseudomonas aeruginosa*, *Candida albicans* or VSV. Our results suggest multiple Toll family members activate the expression of antimicrobial peptides. Our results also provide evidence that Toll and Toll-7 bind multiple Spätzle proteins and differentially affect immune defense against different pathogens in adult male and female flies.

## Introduction

The defense system in *Drosophila melanogaster* can discriminate among various microorganisms and express antimicrobial peptides (AMPs) in response to microbial infection [1,2]. A genetic analysis has revealed that the expression of AMPs is controlled by signaling pathways, such as the spätzle/toll/tube/pelle/cactus gene cassette, which controls for example *drosomycin* expression [3]. Toll (also called Toll-1) was first identified in *Drosophila* as the receptor that regulates the dorsal-ventral patterning in embryonic development [4] and was later found to be involved in the regulation of AMP genes in larvae and adult flies [3].

Since the discovery of *Drosophila* Toll, more than nine Toll-like receptors (TLRs) have been identified in humans and other vertebrate species. Tolls/TLRs are present in all metazoans [5] and mediate important physiological processes, such as inflammation, immune cell regulation, cell survival, and cell proliferation [6-8]. *Drosophila* Toll and mammalian TLRs share a common structural architecture with a conserved ectodomain composed of leucine-rich repeats (LRRs), a single-pass transmembrane domain, and a cytosolic Toll-interlukin-1 receptor homology (TIR) domain, which is also shared by members of the interleukin-1 receptor (IL-1R) family and the intracellular adaptor protein MyD88 [9,10]. However, *Drosophila* Toll and mammalian TLRs show differences in their binding to ligands. *Drosophila* Toll binds to an endogenous protein ligand called Spätzle (Spz, also called Spz-1) [11,12], whereas mammalian TLRs directly recognize various pathogen-associated molecular patterns (PAMPs), such as bacterial lipopolysaccharide (LPS), peptidoglycan (PGN), teichoic acid, flagella and CpG DNA, and viral single-stranded and double-stranded RNAs [13-15].

The *Drosophila* genome encodes nine Tolls [16,17] and six Spätzles [18], and only the functions of the canonical Toll-Spz (also called Toll-1-Spz-1) pathway in development and innate immunity have been well studied. *Drosophila* Toll-2 (18 wheeler, 18W), Toll-5 (Tehao), Toll-8 (Tollo) and Toll-9 may play a role in immunity [16,19-23], Toll-6 and Toll-7 function as neurotrophin receptors and interact genetically with the *Drosophila* neurotrophins DNT1 (Spz-2) and DNT2 (Spz-5) [24]. Toll-7 can recognize *vesicular stomatitis* virus (VSV) and induce antiviral autophagy independently of the canonical Toll pathway [25], but it is not required in the response to anti-VSV infection [26].

In *Drosophila*, Toll activation by Spz is transduced by the adaptor protein MyD88 (dMyD88) via Tube and Pelle kinases to induce the phosphorylation and subsequent degradation of the IκB inhibitor Cactus [27]. Cactus degradation frees the NF-κB factors Dif and/or Dorsal, which translocate(s) to the nucleus to activate the expression of AMP genes [28,29]. Spätzle is synthesized as a pro-protein with an N-terminal prodomain and a C-terminal active cystine knot domain (a.k.a. the cystine-knot family of growth factors), and the activation of pro-Spz requires proteolytic cleavage [30,31]. *Drosophila* pro-Spz (also called pro-Spz-1) is activated by a Spätzle-processing enzyme (SPE) to generate the cystine knot active Spz [32]. Active Spz dimers bind to two Toll receptors to trigger the downstream signaling pathway [12,33,34], but a direct interaction between different Toll and other Spz proteins has not yet been reported.

The cystine knot domains of Spz-2 to Spz-6 can be predicted based on their amino acid sequences, and these are located in the middle region of pro-Spz-2 and at the C-termini of pro-Spz-3 to pro-Spz-6, respectively (see Fig EV1A). However, the functions of Spz-2 to Spz-6 as ligands for the activation of the Toll pathway in *Drosophila* innate immunity have not yet been reported.

In this study, we report that in addition to Toll, *Drosophila* Toll-7 also strongly activated the *drosomycin* promoter. More importantly, we showed that Toll and Toll-7 interacted with Spz, Spz-2 and Spz-5, and multiple pairs of Spz proteins with Toll or Toll-7 activated the *drosomycin* promoter. Furthermore, Toll and Toll-7 both recognized VSV by binding to the VSV glycoprotein; Toll and Toll-7 differentially affected defense responses in adult male and female flies after systemic infection by *Enterococcus faecalis*, *Pseudomonas aeruginosa*, *Candida albicans* or VSV. Our results suggest multiple Toll family members activate the expression of antimicrobial peptides. Our results also provide evidence that Toll and Toll-7 bind multiple Spz proteins and differentially affect immune defense against different pathogens in adult male and female flies.

## Results

### TIR domains of *Drosophila* Tolls activate the *drosomycin* but not the *diptericin* promoter in S2 cells

The *Drosophila* Toll-Spätzle (Toll-1-Spz-1) pathway is activated after the binding of the active Spz dimer to two Toll receptors. This binding triggers dimerization of the intracellular TIR domains and their subsequent interaction with the adaptor protein dMyD88 [35,36] to relay intracellular signals, which induce the translocation of the NF-κB factors Dorsal/Dif into the nucleus to activate AMP genes, such as *drosomycin* [28,29].

To test whether *Drosophila* Tolls serve as functional receptors in the activation of AMP genes, the TIR domains of *Drosophila* Toll to Toll-9 and *M. sexta* Toll [37], all contain only the intracellular domains (without the single-pass transmembrane domains), were overexpressed in S2 cells, and the activation of the *drosomycin* and *diptericin* promoters by TIR domains was assessed through dual luciferase assays because the overexpression of TIR domains can lead to the formation of TIR dimers/oligomers, which can recruit dMyD88 to trigger the downstream intracellular signaling pathway. Western blot results showed that the TIR domains of all ten Tolls were expressed in S2 cells (Fig 1A and B). In addition, dual luciferase assays showed that overexpression of the TIR domains from the nine *Drosophila* Tolls and *M. sexta* Toll activated the *drosomycin* promoter to certain extents, and increased promoter activity was observed with the TIR domains of Toll, Toll-7 and *M. sexta* Toll, but overexpression of the TIR domains from all ten Tolls did not activate the *diptericin* promoter (Fig 1C). These results suggested that all nine *Drosophila* Tolls play a role in immune signaling pathways and that Toll and Toll-7 might play a major role in these pathways. We focused on Toll and Toll-7 in our subsequent study as they can strongly activated the *drosomycin* promoter.

**Figure 1.**
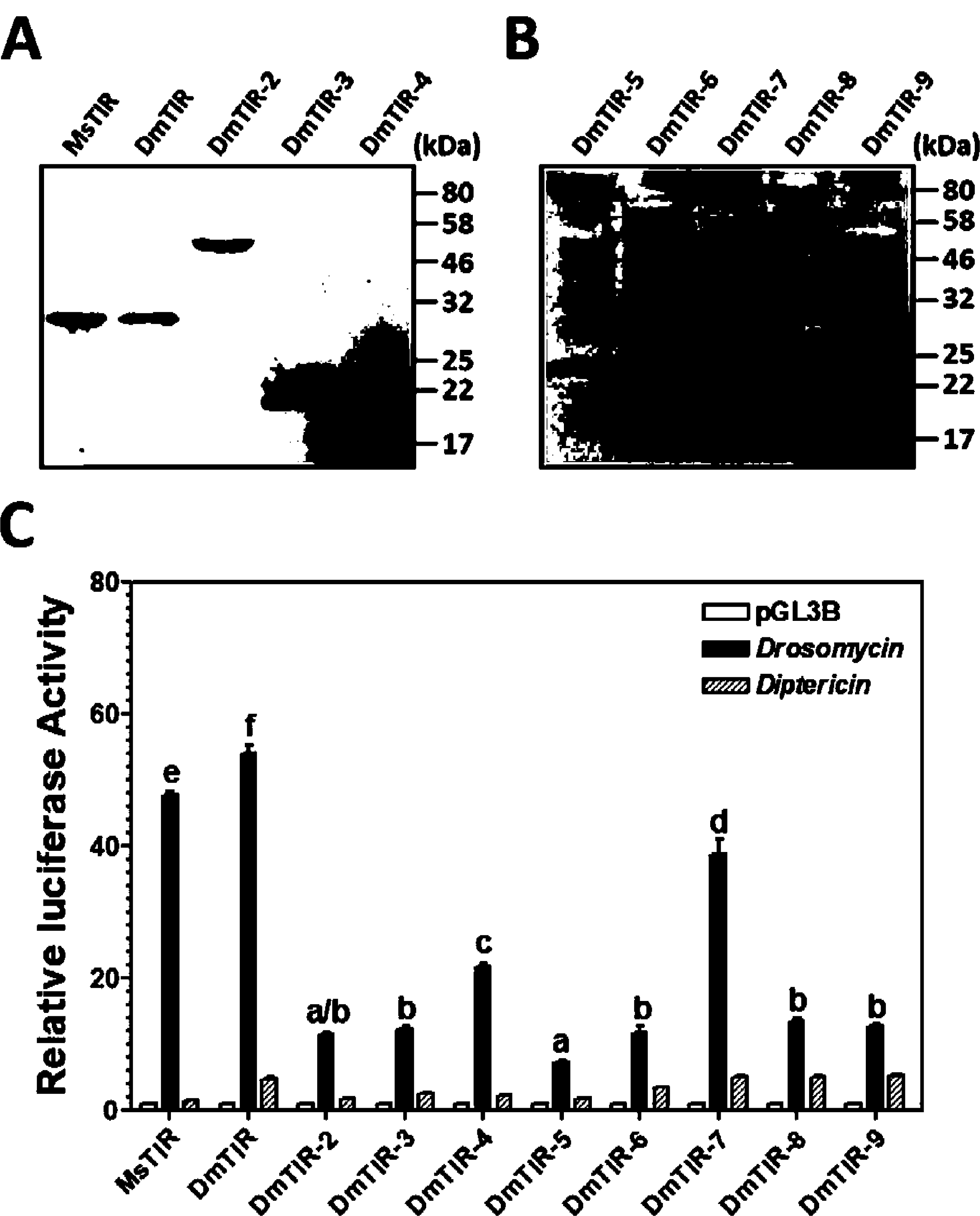
Activation of the *drosomycin* promoter through the overexpression of TIR domains. A, B Expression of V5-tagged TIR domains from *Drosophila* Toll to Toll-9 and *M. sexta* Toll in S2 cells detected by anti-V5 antibody. C Relative luciferase activity of *drosomycin* and *diptericin* promoter reporters in S2 cells overexpressing TIR domains. Data information: In C, the graph shows mean ± SEM, n=3; identical letters show a non-significant difference (*p* > 0.05), whereas different letters indicate a significant difference (*p* < 0.05), one-way ANOVA followed by Tukey’s multiple comparison test.

### The ectodomains of Toll and Toll-7 interact with multiple Spätzle proteins

The cystine knot domains of *Drosophila* Spz-2 to Spz-6 were predicted based on their amino acid sequences (Fig EV1A). Among the six Spz proteins, Spz, Spz-2 and Spz-5 share high similarities (Spz shares 63% and 71% similarities with Spz-2 and Spz-5, respectively, and Spz-2 shares 62% similarity with Spz-5) (Fig EV1B), and these three proteins are phylogenetically more closely related. Through co-immunoprecipitation (Co-IP) assays, we previously showed that Toll receptors interact with the active cystine knot domains of Spätzles but not the full-length pro-Spätzles [37]. We expressed recombinant Toll^ecto^, Toll-7^ecto^ (ectodomains), Toll and Toll-7 (full-length receptors) as well as the cystine knot domains of Spz to Spz-6 in S2 cells. Toll^ecto^, Toll-7^ecto^ and the six Spz proteins were detected in both the cell culture media and the cell lysates, whereas full-length Toll and Toll-7 were detected only in the cell lysates and not in the cell culture media (Fig EV2). Co-IP assays showed that Toll^ecto^ interacted with Spz, Spz-2 and Spz-5 but not with Spz-3, Spz-4 or Spz-6 (Fig 2A-D), whereas Toll-7^ecto^ interacted with Spz, Spz-2, Spz-5 and Spz-6 but not with Spz-3 or Spz-4 (Fig 2E-H). These findings suggested that Toll and Toll-7 can bind to multiple Spz ligands.

**Figure 2.**
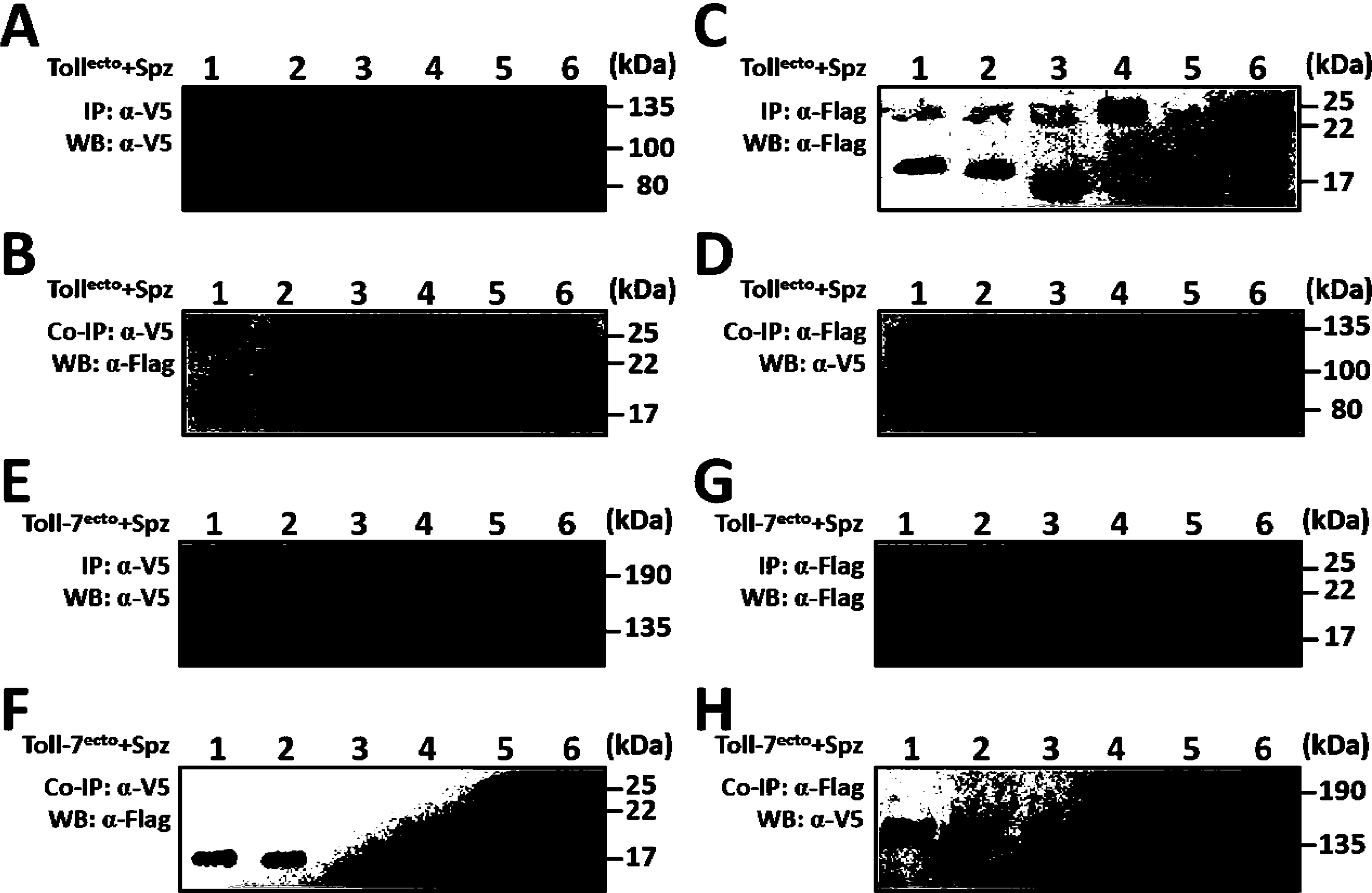
Interaction of Toll and Toll-7 ectodomains with Spz proteins. Recombinant V5-tagged ectodomains of Toll and Toll-7 (Toll^ecto^ and Toll-7^ecto^) and the Flag-tagged active cystine knot domains of Spz to Spz-6 were overexpressed in S2 cells separately, and cell lysates from Toll^ecto^ (or Toll-7^ecto^) and one of the six Spz proteins were mixed for co-immunoprecipitation (Co-IP) assays. Lanes 1–6 were Spz to Spz-6 mixed with Toll^ecto^ or Toll-7^ecto^. A, E Anti-V5 antibody precipitated Toll^ecto^ (A) and Toll-7^ecto^ (E) proteins, and these were detected with anti-V5 monoclonal antibody. B, F Spz proteins co-immunoprecipitated with Toll^ecto^ (B) and Toll-7^ecto^ (F) and were detected with anti-Flag monoclonal antibody. C, G Anti-Flag antibody precipitated Spz proteins, and these were detected with anti-Flag antibody. D, H Toll^ecto^ (D) and Toll-7^ecto^ (H) co-immunoprecipitated with Spz proteins and were detected with anti-V5 antibody.

### Multiple Toll-Spz and Toll-7-Spz pairs activate the *drosomycin* promoter in S2 cells

To determine whether multiple pairs of Spz proteins with Toll or Toll-7 can trigger signaling pathways, dual luciferase assays were performed. The co-expression of Toll with Spz, Spz-2 and Spz-5 activated the *drosomycin* promoter, and the highest activity was obtained with Toll-Spz, followed by Toll-Spz-2 and Toll-Spz-5 (Fig 3A). In addition, the co-expression of Toll-7 with Spz, Spz-2 and Spz-5 also activated the *drosomycin* promoter, and the highest activity was observed with Toll-7-Spz-2, followed by Toll-7-Spz-5 and Toll-7-Spz (Fig 3B). These results are consistent with those obtained with the interaction of Toll and Toll-7 with Spz, Spz-2 and Spz-5 (Fig 2). We also confirmed that the overexpression of Tolls (Toll, Toll-2 and Toll-7) or Spz proteins (Spz, Spz-2 and Spz-5) alone, and the co-expression of Toll-2 and Spz (non-functional pair) did not activate the *drosomycin* promoter, and that only the co-expression of the correct pairs of Toll and Spz proteins (Toll-Spz, Toll-7-Spz and Toll-7-Spz-5) activated the *drosomycin* promoter (Fig 3C). The expression or co-expression of all these proteins did not activate the *diptericin* promoter (Fig 3).

**Figure 3.**
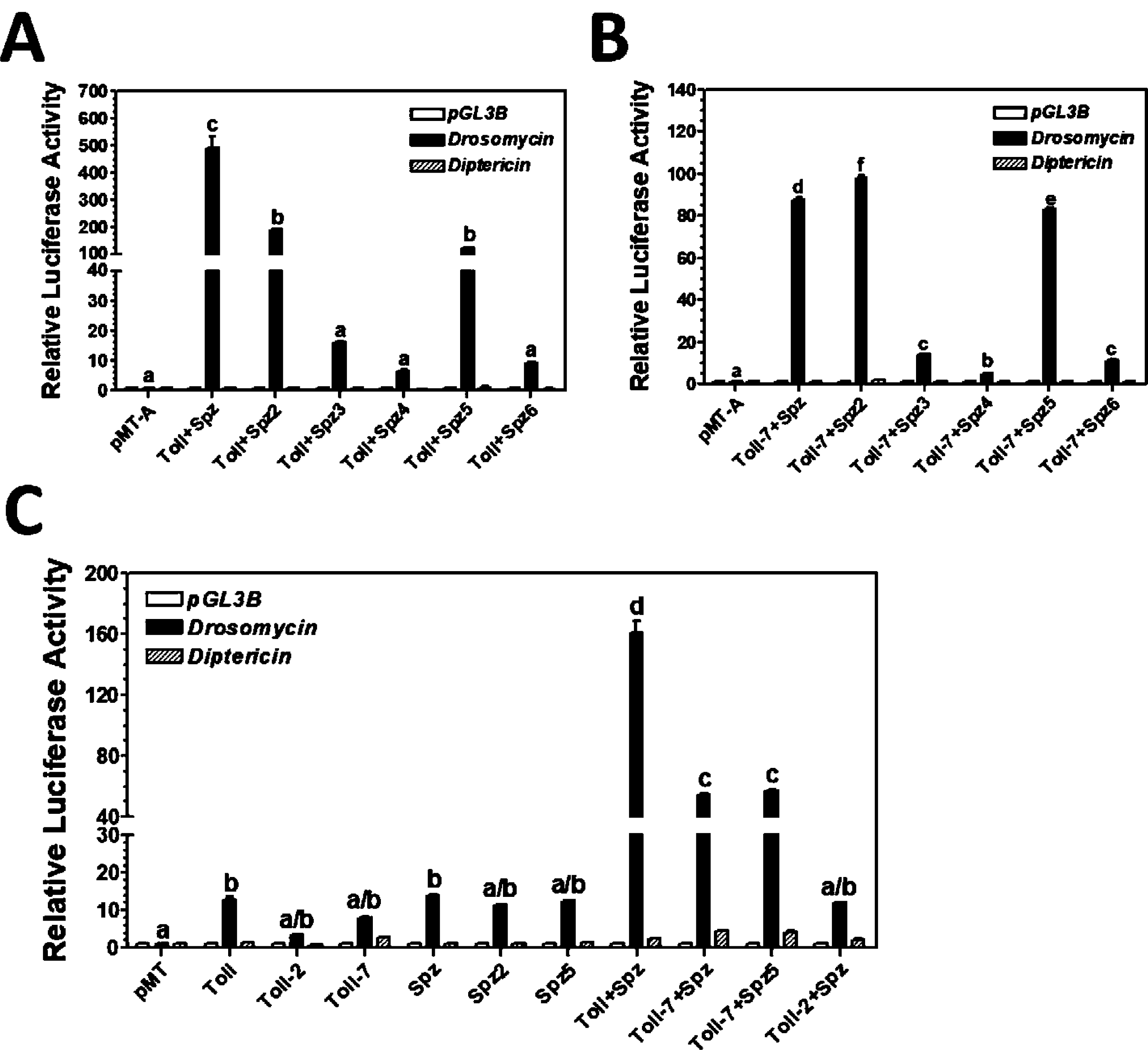
Activation of the *drosomycin* promoter by multiple pairs of Toll-Spz and Toll-7-Spz. A-C The relative luciferase activity of the *drosomycin* or *diptericin* promoter reporter in S2 cells overexpressing full-length Toll and one of the six Spz proteins (A), overexpressing Toll-7 with one of the six Spz proteins (B), or overexpressing individual Toll or Spz proteins or different pairs of Toll and Spz proteins (C) was determined using a Dual-Luciferase® Reporter Assay System. Data information: In (A-C), the graphs show mean ± SEM, n=3; identical letters show a non-significant difference (*p* > 0.05), whereas different letters indicate a significant difference (*p* < 0.05), one-way ANOVA followed by Tukey’s multiple comparison test.

### Ectodomains of Toll and Toll-7 recognize VSV, and VSV infection activates AMP gene promoters

Toll-7 can recognize VSV but is not involved in the anti-VSV response [25,26]. We first determined whether Toll can also recognize VSV. Toll^ecto^ and Toll-7^ecto^ were overexpressed in S2 cells and secreted into the cell culture media, and the VSV glycoprotein (VSV-G) was detected in the virus-infected DMEM cell culture media (Fig 4A). When the cell culture media containing Toll^ecto^ and Toll-7^ecto^ were mixed with VSV virions and the V5-tagged ectodomains were pulled down by anti-V5 antibody, Toll^ecto^ and Toll-7^ecto^ were detected in the immunoprecipitated proteins (Fig 4B), and VSV-G was also detected in the Co-IP proteins (Fig 4C), indicating that both Toll^ecto^ and Toll-7^ecto^ recognize VSV through interaction with VSV-G. When stable S2 cell lines expressing full-length Toll and Toll-7 were transfected with AMP gene promoter reporters, the activity of the AMP gene promoters, including *drosomycin* and *attacin* promoters, were significantly activated by VSV infection (Fig 4D and E), suggesting that the recognition of VSV by Toll and Toll-7 can activate AMP genes.

**Figure 4.**
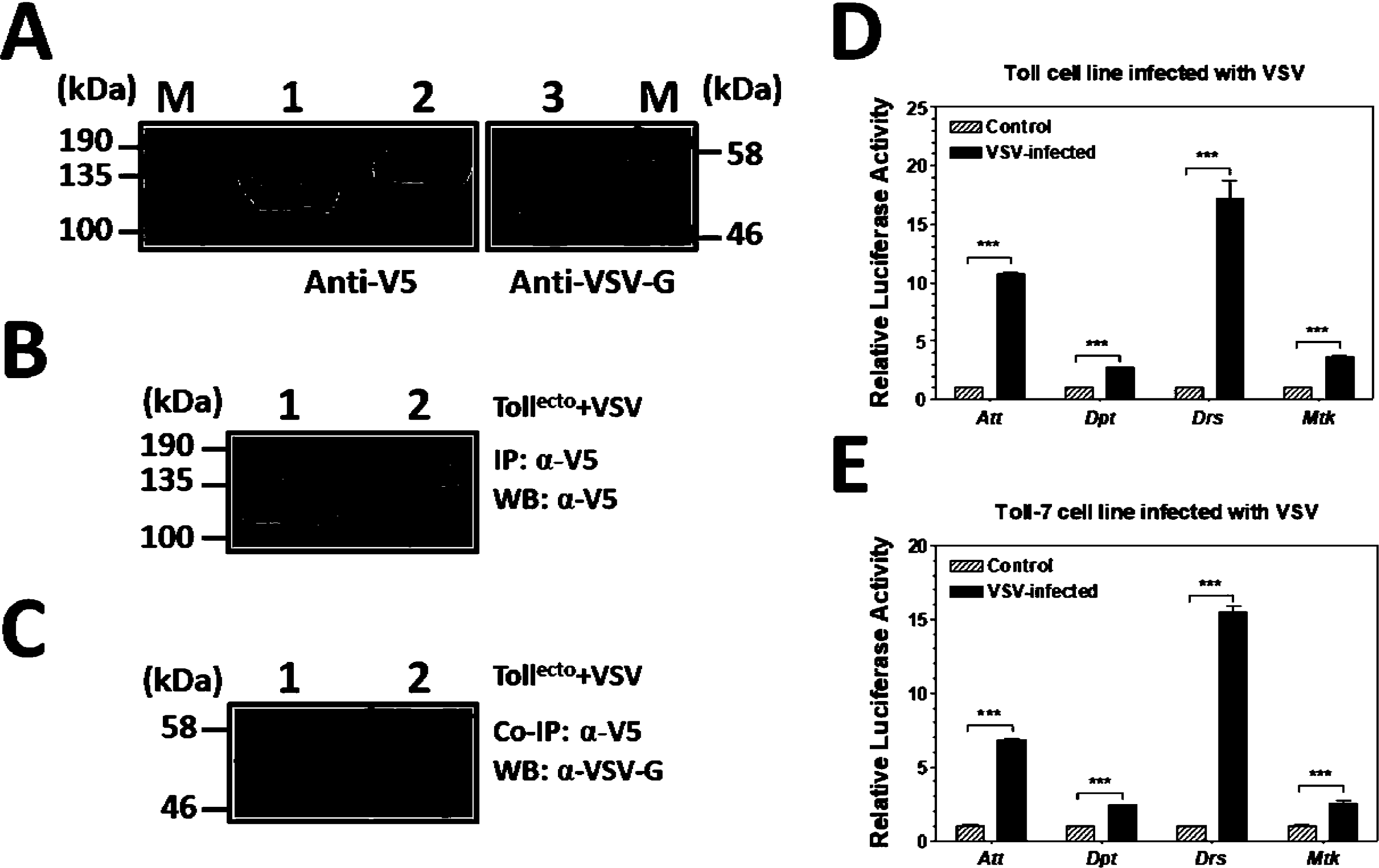
Interaction of Toll and Toll-7 ectodomains with VSV and activation of AMP gene promoters by VSV infection. A Expression of Toll^ecto^ (lane-1) and Toll-7^ecto^ (lane-2) in S2 cell culture media detected by anti-V5 antibody, and detection of VSV glycoprotein (VSV-G) (lane 3) in the VSV-infected cell culture media with anti-VSV-G antibody. B, C Interaction of Toll^ecto^ and Toll-7^ecto^ with VSV-G. V5-tagged Toll^ecto^ and Toll-7^ecto^ were mixed with VSV virions, and proteins were pulled down with anti-V5 antibody. Immunoprecipitated Toll^ecto^ (lane-1) and Toll-7^ecto^ (lane-2) were detected with anti-V5 antibody (B), and co-immunoprecipitated VSV-G protein was detected with anti-VSV-G antibody (C). D, E Activation of AMP gene promoter reporters in Toll and Toll-7 cell lines by VSV infection. Stable S2 cell lines expressing recombinant full-length Toll and Toll-7 were transfected with AMP gene promoter reporters and then infected with VSV. The relative luciferase activity was determined using a Dual-Luciferase^®^ Reporter Assay System. Data information: In (D, E), the graphs show mean ± SEM, n=3; *** *p* < 0.001, unpaired t-test.

### Toll and Toll-7 play differential roles in male and female flies to defend against bacterial, fungal and VSV infection

To verify the functions of Toll and Toll-7 in *Drosophila* immunity, wild-type (*w*^*1118*^) flies, various Toll mutants (*Tlr*^*632*^*/Tl*^*I-RXA*^ and *Tl*^*I-RXA*^/*TM6B*) and Toll-7 mutants (*Toll-7*^*g1-5*^*/CyO* and *Df(2R)BSC22*/*Toll-7*^*g1-5*^) were infected with the pathogenic bacteria *E. faecalis* and *P. aeruginosa*, *C. albicans*, or VSV-GFP, and the cumulative survival of the flies was recorded. In *w*^*1118*^ flies, *Toll* was expressed at a slightly lower level than *Toll-7* in males but at a significantly higher level than *Toll-7* in females (Fig EV4A). Therefore, septic infection assays were separately performed in male and female flies. Compared with *w*^*1118*^ flies, all four mutant males and the two *Toll* mutant females were significantly more susceptible to *E. faecalis* infection (Fig 5A and E). In addition, the two *Toll* mutant males were significantly more susceptible to *E. faecalis* infection than the two *Toll-7* mutant males (Fig 5A), and the two *Toll-7* mutant males were significantly more susceptible to *P. aeruginosa* infection than the two *Toll* mutant males (Fig 5B). Moreover, all four mutant males and the two *Toll* mutant females were significantly more susceptible to *C. albicans* infection (Fig 5C and G), and the *Df*/*Toll-7*^*g1-5*^ and *Tlr*^*632*^*/Tl*^*I-RXA*^ males and all four mutant females were significantly more susceptible to VSV-GFP infection (Fig 5D and H). Additionally, the two *Toll* mutant females were significantly more susceptible to VSV-GFP infection than the two *Toll-7* mutant females (Fig 5H). The detection of *gfp* transcript expression in the VSV-GFP infected flies revealed no significant differences among the *w*^*1118*^, *Df*/*Toll-7*^*g1-5*^ and *Tlr*^*632*^*/Tl*^*I-RXA*^ flies at days 1, 5 and 10 post-infection (Fig EV3), indicating that the VSV-GFP titer remains at a similar level in these flies even 10 days after infection. Taken together, these results suggest that Toll and Toll-7 are required for both *Drosophila* males and females in defense against microbial infections. Whether Toll or Toll-7 plays a major/minor role may be related to the differential expression of *Toll* and *Toll-7* transcripts in *w*^*1118*^ males and females (Fig EV4A) and the induced/reduced expression of *Toll*, *Toll-7* (Fig EV4B-I) and AMP genes (Fig EV5) in mutant flies after microbial infection. For example, *Toll-7* was expressed at a significantly lower level in all four mutant females than in the *w*^*1118*^ females after microbial infection (Fig EV4F-I), but *Toll* was induced in the two *Toll* mutant females after *E. faecalis* infection (Fig EV4F) and in the *Toll-7*^*g1-5*^*/CyO* females after *P. aeruginosa* (Fig EV4G) and *C. albicans* infection (Fig EV4H). *Toll-7* was significantly up-regulated in the two *Toll* mutant males after *P. aeruginosa* (Fig EV4C) and *C. albicans* (Fig EV4D) infection and in the *Tl*^*I-RXA*^/*TM6B* males after *E. faecalis* infection (Fig EV4B) compared with *w*^*1118*^ males. *Drosomycin* was expressed at a significantly lower level in all four mutant flies compared with the *w*^*1118*^ flies after *E. faecalis* infection (Fig EV5A and E) but was significantly up-regulated in the *Toll-7*^*g1-5*^*/CyO* and the two *Toll* mutant flies compared with the *w*^*1118*^ flies after VSV-GFP infection (Fig EV5D and H).

**Figure 5.**
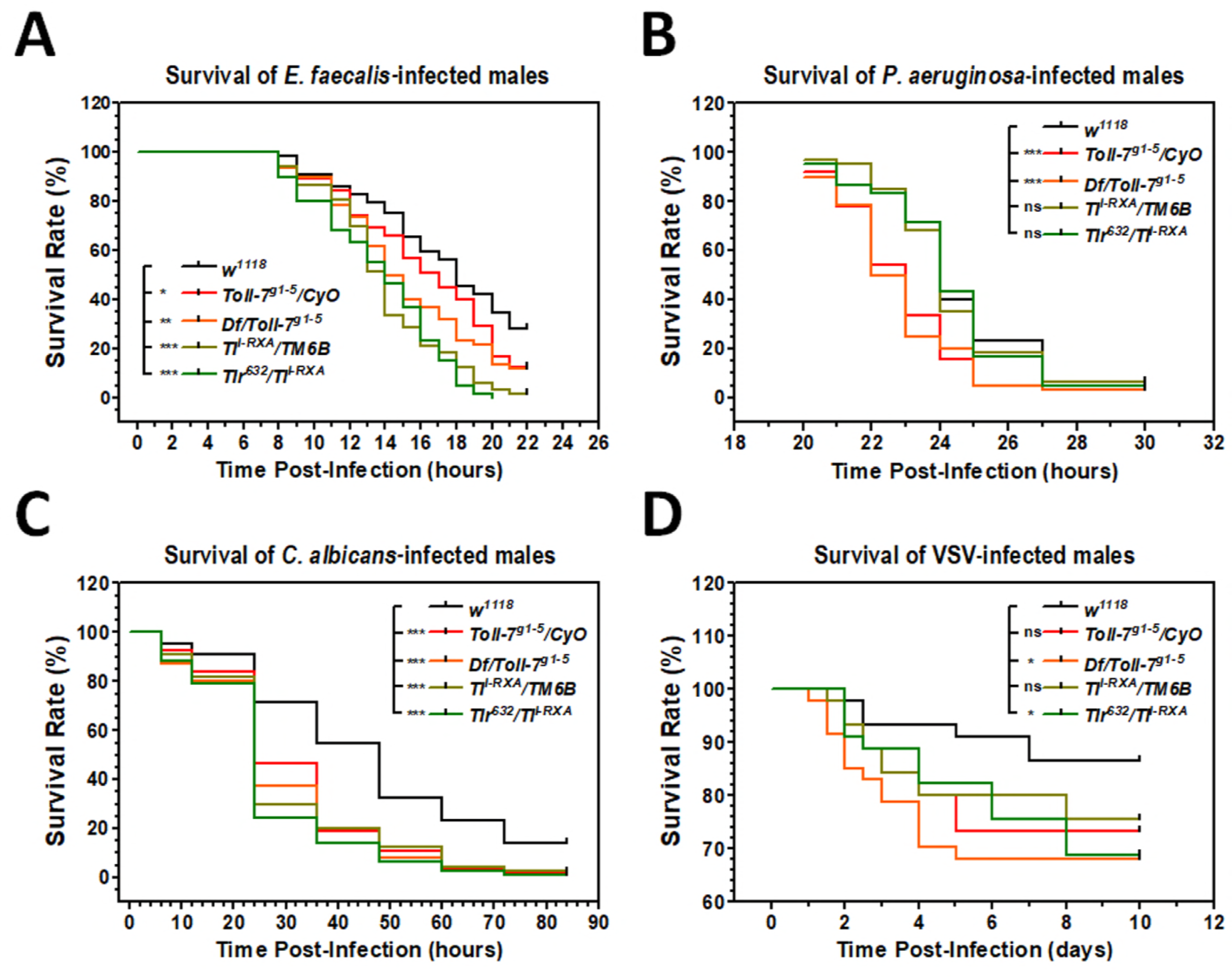
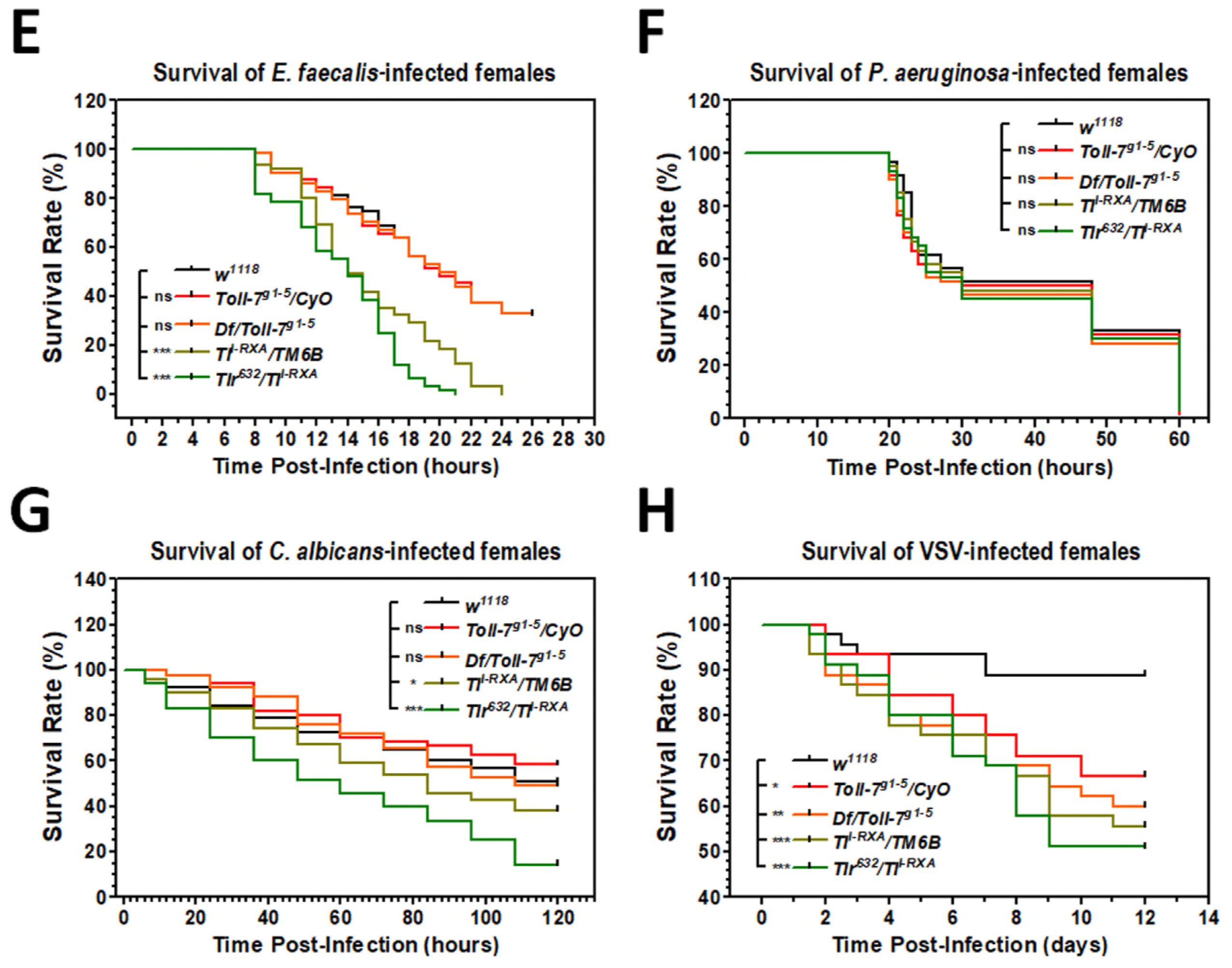
Toll and Toll-7 confer resistance to microbial infection in adult flies. A-H *w*^*1118*^, *Toll-7*^*g1-5*^*/CyO*, *Df(2R)BSC22/Toll-7*^*g1-5*^, *T1*^*I-RXA*^/*TM6B* and *Tlr*^*632*^*/Tl*^*I-RXA*^ mutant males (A-D) and females (E-H) were infected with *E. faecalis*, *P. aeruginosa*, *C. albicans* and VSV-GFP, and the cumulative survival of the flies was recorded. Data information: In (A-H), * *p* < 0.05, ** *p* < 0.01 and *** *p* < 0.001; ‘ns’ for not significant difference when compared with *w*^*1118*^, unpaired t-test.

## Discussion

Among the nine *D. melanogaster* Tolls [16,17] and six Spätzles [18], only Toll and Spz have been well studied [3]. Although there are reports about Toll-2, Toll-5, Toll-8 and Toll-9 in activation of AMPs expression [19-23,38,39], there has been no reports about Spz-2 to Spz-6 as ligands for Tolls in immune signaling pathways. We report here that Spz, Spz-2 and Spz-5 formed multiple Spz-Toll pairs with Toll and Toll-7, and multiple Spz-Toll and Spz-Toll-7 complexes activated *drosomycin* promoter, suggesting multiple *Drosophila* Toll-Spz pathways in regulating the expression of antimicrobial peptide genes.

In 2000, Tauszig et al. [16] compared eight *Drosophila* Tolls (Toll to Toll-8) in activation of AMP gene promoters. They expressed chimeric Tolls, in which each of the TIR domain from Toll-2 to Toll-8 was fused to the truncated extracellular domain of Toll, as the truncated Toll (Toll^ΔLRR^) that has the leucine-rich repeats (LRRs) deleted is an active receptor independence of Spz ligand [40]. Among the five chimeric Tolls (Toll-2, Toll-5 to Toll-8) that were expressed in S2 cells, only expression of chimeric Toll-5 activated *drosomycin* promoter activity to about 25% of that activated by Toll^ΔLRR^, and expression of all five chimeric Tolls and Toll^ΔLRR^ did not activate *diptericin*, *defensin* and *cecropin* promoters [16]. We expressed only the TIR domains (without the extracellular and transmembrane domains) of Toll to Toll-9 and found that expression of all nine TIRs activated *drosomycin* promoter to some extents, with high activity observed with TIR of Toll, followed by TIRs of Toll-7 and Toll-4. As an active receptor without ligand binding, Toll^ΔLRR^ must be able to form dimers/oligomers. It is possible that except chimeric Toll-5, the other four chimeric Tolls (Toll-2, Toll-6 to Toll-8) could not form stable dimers/oligomers, while free TIRs expressed in the cytoplasm in our study may easily form dimers/oligomers. Expression of all TIRs did not activate *diptericin* promoter, a result consistent with that from expression of chimeric Tolls [16].

Toll-6 and Toll-7 can function as neurotrophin receptors in the development of the central nervous system in *Drosophila* and may interact with Spz-2 and Spz-5, respectively [24]. We showed that Toll-7 interacted with Spz-6, but that co-expression of Toll-7 with Spz-6 did not activate the *drosomycin* promoter, suggesting that Toll-7-Spz-6 may play a role in development. Multiple pairs of Spz-Toll-7 in activation of *drosomycin* promoter as well as Toll-7 mutant male and female flies were significantly more susceptible to microbial infection suggest that Toll-7, like Toll, play roles in both innate immunity and development.

Toll-7 can recognize *vesicular stomatitis* virus (VSV) to induce anti-viral autophagy response [25], but it is not required for anti-VSV infection [26]. We confirmed that Toll-7 recognized VSV through binding to VSV glycoprotein. In addition, we showed that Toll also bound VSV glycoprotein, and binding of VSV to Toll and Toll-7 activated AMP gene promoters. Septic infections with different pathogens in *w*^*1118*^, *Toll* and *Toll-7* mutant flies suggest that Toll and Toll-7 differentially affect adult male and female flies in defense against infection by *E. faecalis*, *P. aeruginosa*, *C. albicans* and VSV. As Toll and Toll-7 both play a role in defense against microbial infection, the survival of male and female flies after septic infections depends on the overall effect of differential expression levels of *Toll*, *Toll-7* and AMP genes. The titers of VSV-GFP (determined by the *gfp* transcript) in *w*^*1118*^, *Toll* and *Toll-7* mutant flies maintained at the similar levels at day 1, 5 and 10 post-infection in each fly line as well as among the three fly lines, indicating that Toll and Toll-7, though play roles in defense against VSV infection, may not play a role in restricting VSV replication.

We demonstrated the existence of multiple Toll-Spz pathways in *Drosophila* innate immunity, which raises more questions that need to be answered. What are the functions of different Toll-Spz and Toll-7-Spz pairs in *Drosophila* immunity? How pro-Spz-2 and pro-Spz-5 are processed/activated (particularly pro-Spz-2, which requires two proteolytic cleavages)? Are Spz-2 and Spz-5 processed by SPE or/and other unidentified proteases? What is the function of Toll-7-Spz-6? Future research will focus on answering some of these questions.

## Materials and Methods

### Fly stocks

Wild-type *w*^*1118*^ flies were obtained from the laboratory of Dr. Leonard Dobens (School of Biological Sciences, University of Missouri – Kansas City, Missouri, USA). The *Toll-7*^*g1-5*^*/CyO* mutant line was a gift from Dr. Yashimasa Yagi (Division of Biological Science, Nagoya University, Nagoya, Japan), and the *Toll-7*^*g1-5*^ mutant line was created by homologous recombination of an ends-in knockout system followed by hs-ICreI treatment to generate a *Toll-7* knockout line with a point mutation [17]. The *Toll-7*^*g1-5*^ line was balanced over CyO to obtain the *Toll-7*^*g1-5*^*/CyO* mutant line, and heterozygotes were screened based on the existence of curled wings. The *Tlr*^*632*^*/Tl*^*I-RXA*^ and *Tl*^*I-RXA*^/*TM6B* Toll (also called Toll-1) mutant lines were obtained from the laboratory of Dr. Kontoyiannis (Department of Infectious Diseases, University of Texas M. D. Anderson Cancer Center, Houston, Texas, USA) [41]. *Tlr*^*632*^/*Tl*^*I-RXA*^ flies were generated by crossing *Tlr*^*632*^/*TM6B* and *Tl*^*I-RXA*^/*TM6B Toll*-deficient flies. *Tlr*^*632*^ is a thermosensitive loss-of-function allele with a strong phenotype at 29^°^C; thus, these flies were maintained at 29^°^C during infection. Both the *Tlr*^*632*^ and *Tl*^*I-RXA*^ mutant lines were balanced over *TM6B* and were recognized by multiple hair-type bristle in the upper lateral thorax/torso. The *Df(2R)BSC22/SM6a* line (stock # 6647) was purchased from Bloomington Stock Center (Indiana, USA); in this line, the 56D7 – 56F12 chromosome segment was deleted by exploiting hybrid element insertion (HEI) and resolution, and this line was later balanced over *SM6a* to obtain flies that can be recognized by curly wings. We generated *Df(2R)BSC22*/*Toll-7*^*g1-5*^ flies by crossing *Toll-7*^*g1-5*^*/CyO* and *Df(2R)BSC22/SM6a* flies, which uncovers the *Toll-7* locus to obtain *Toll-7* mutants that can be screened by the presence of curly wings. All the flies were cultured on corn-meal diet [42] and transferred to fresh food at least 24 h prior to injection/infection.

### Gene cloning

The nine *Drosophila* Toll clones are available in Dr. Y. Tony Ip laboratory at the University of Massachusetts Medical School, Worcester, MA, USA [17]. All these clones were genomic DNAs cloned in the pAC5.1-A vectors and might thus contain introns. For this study, we cloned Toll cDNA using the total RNA from *Drosophila* adult females as the template and Toll-7 cDNA using the pAC5.1-A clone, which did not contain any introns, as the template. All nine *Drosophila* Toll TIR domains and *M. sexta* Toll TIR domain [37], the ectodomains of Toll and Toll-7 and full-length Toll and Toll-7 were amplified by PCR using the forward and reverse primers listed in Table 1 and cloned into the pMT/BiP/V5-His A vector (V413020, Invitrogen) for expression of the recombinant proteins with a V5-tag at the C-terminus. Active Spz to Spz-6 proteins (Fig EV1A) were amplified by PCR and cloned into a modified pMT/Bip A vector [37] for expression of the recombinant Spz proteins with a Flag-tag at the N-terminus. The PCR reactions were performed with the following conditions: 94^°^C for 3 min, 35 cycles of 94^°^C for 30 s, Tm-5^°^C for 30 s, 72^°^C for 30 s to 4 min, and final extension at 72^°^C for 10 min. The PCR products were recovered using an Agarose Gel Electrophoresis-Wizard® SV Gel and PCR Clean-Up System (A9285, Promega) and then subcloned into the T-Easy vector (A1360, Promega). Recombinant plasmid DNAs were purified using a PureYield™ Plasmid Miniprep System (A1222, Promega) according to the manufacturer’s instructions and digested with respective restriction enzymes, and DNA fragments were recovered and inserted into the pMT/BiP/V5-His A or modified pMT/Bip A vector using T4 DNA ligase (M0202L, NEB). The recombinant expression plasmids were then purified and sequenced in the sequencing facility at University of Missouri – Columbia for further experiments.

**Table 1.**
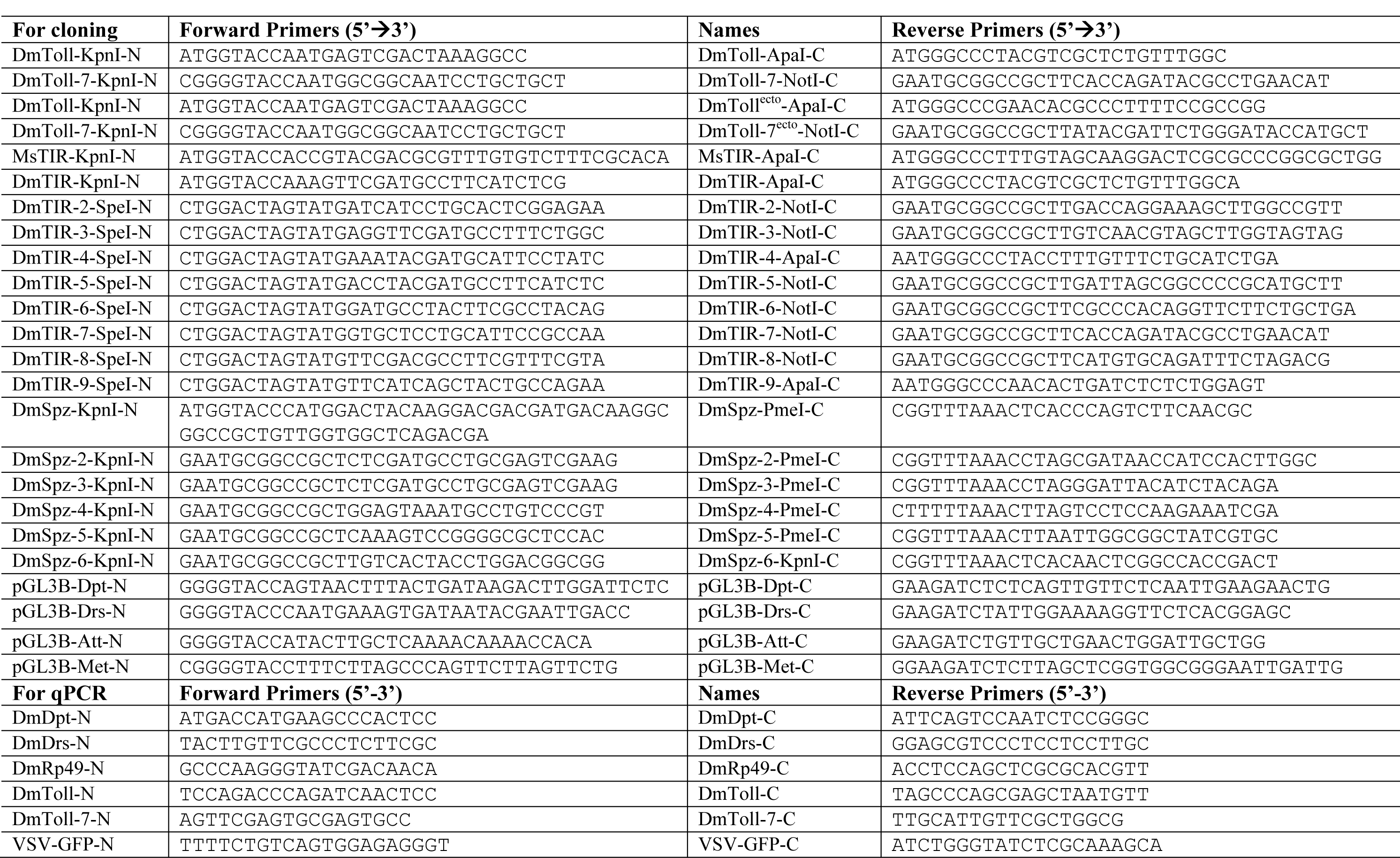
Primers used in this study.

### *Vesicular stomatitis* virus stock culture

*Vesicular stomatitis* virus (VSV) expressing green fluorescent protein (VSV-GFP), in which GFP is inserted between the 3’ leader and N gene [43], was obtained from the laboratory of Dr. Whelan (Harvard Medical School, Boston, Massachusetts, USA) [44]. VSV-GFP was cultured and maintained in HEK293 cells in DMEM medium supplemented with 10% heat-inactivated fetal bovine serum (#10082063, Invitrogen) and 1% penicillin-streptomycin solution (G6784, Sigma-Aldrich). The viral titer was measured by plaque assay using HEK293 cells [45]. For infection assays with *Drosophila* S2 cells, 10,000 pfu/ml VSV-GFP was used, and for the infection assays with adult flies, 10,000 pfu of VSV-GFP (in 50 nl of PBS) were injected into each fly.

### Infection assays

*Drosophila* adult males and females (5-7 days of age) in a batch of 20-30 flies were infected with the Gram-positive bacterium *Enterococcus faecalis V583* (a gift from Dr. Michael Gilmore, Harvard Medical School, Boston, Massachusetts, USA), the Gram-negative bacterium *Pseudomonas aeruginosa PA-14* (a gift from Dr. Kalai Mathee, Florida International University, Florida, USA), the fungus *Candida albicans* (a gift from Dr. Theodore White, School of Biological Sciences at the University of Missouri – Kansas City, Missouri, USA), or VSV-GFP. Briefly, overnight bacterial and fungal cultures were diluted to OD_600_ = 0.2 and OD_600_ = 0.5, respectively, washed with phosphate buffered saline (PBS, pH 7.4) and resuspended in PBS for injection. Flies were anesthetized with CO_2_ (for no longer than 15 min at a time), and 50 nl of diluted *E. faecalis V583*, *P. aeruginosa PA-14*, *C. albicans*, or VSV-GFP (10,000 pfu/50 nl) was injected into each fly at the left intra-thoracic region using a Drummond nanoinjector and pulled glass capillary needles. These flies were maintained in clean bottles with fresh cornmeal diet, and the diet was changed every day throughout the course of the experiment. The flies that died within 3 h of injection were excluded from the study due to death by injury. The flies were monitored every hour (or every day in the VSV-GFP infection assay), and the numbers of dead males and females were recorded. The accumulation of VSV-GFP RNA in the *w*^*1118*^, *Df(2R)BSC22*/*Toll-7*^*g1-5*^ and *Tlr*^*632*^*/Tl*^*I-RXA*^ flies infected with VSV-GFP at day 1, 5 and 10 post-infection was detected by real-time PCR analysis of the *gfp* transcript using primers for GFP (Table 1).

For the infection of *Drosophila* S2 cells with VSV-GFP, S2 cells were grown and maintained in insect cell culture media (SH30610.02, HyClone) supplemented with 10% heat-inactivated fetal bovine serum, 1% penicillin-streptomycin solution and 1% Gibco L-Glutamine (25030081, Thermo Fisher Scientific, complete growth medium). VSV-GFP was cultured in HEK293 cells as described above, and the replication of VSV-GFP was detected by Western blot analysis using anti-VSV-G antibody. A stable S2 cell line expressing either full-length Toll or Toll-7 receptor (described below) was transiently transfected with pGL3B or the pGL3B-*attacin*, pGL3B*-diptericin*, pGL3B-*drosomycin* or pGL3B*-metchnikowin* AMP gene promoter individually using Gencarrier-2 (#31-00110, Epoch Biolabs). Forty-eight hours after protein expression, the S2 cells were infected with 10,000 pfu/ml VSV-GFP for 24 h and processed for dual luciferase assay (see below).

### Transient transfection and establishment of stable S2 cell lines

Transient transfection experiments and the establishment of stable S2 cell lines expressing full-length Toll and Toll-7 were performed as described previously [37]. For transient transfections, S2 cells were seeded overnight in complete growth medium (see above), washed with serum-free medium (SH30278.01, HyClone), and transiently transfected using GenCarrier-2™ transfection reagent (#31-00110, Epoch Biolabs) according to the manufacturer’s instructions. DES®–Inducible/Secreted Kit with pCoBlast (K5130-01, Invitrogen) was used for the establishment of stable S2 cell lines. To select stable S2 cells expressing recombinant Toll and Toll-7, pCoBlast (Invitrogen) was co-transfected with recombinant pMT/BiP/V5-His A vectors. Forty-eight hours after transfection, S2 cells were centrifuged and resuspended in the complete growth medium containing 25 μg/ml Blasticidin S hydrochloride (No.15205, Sigma-Aldrich). Resistant colonies appeared one week later.

### Dual luciferase assays

Dual luciferase assays were performed as described previously [46]. S2 cells were plated in 24-well culture plates (3 × 10^5^ cells/well) overnight in the complete growth medium, washed with serum-free medium, and then transiently co-transfected with recombinant pMT/BiP/V5-His A expression plasmid (500 ng), pGL3B, pGL3B*-drosomycin* or pGL3B*-diptericin* firefly luciferase reporter plasmid (250 ng), or Renilla luciferase reporter plasmid (25 ng) (as an internal standard) (pRL-TK, Promega) with Gencarrier-2. After overnight transfection, serum-free medium was replaced with the complete growth medium containing copper sulfate (to a final concentration of 500 μM) for protein expression, and 36 h after protein expression, the firefly luciferase and Renilla luciferase activities were measured using the Dual-Luciferase Reporter Assay System (E1980, Promega) with a GloMax® Multi Microplate Luminometer (Promega). The relative luciferase activity (RLA) was obtained as the ratio of firefly luciferase activity to Renilla luciferase activity. The RLA obtained for S2 cells co-transfected with empty pMT/BiP/V5-His A and pGL3B (empty reporter vector) plasmids was used as the calibrator. These experiments were repeated at least three times (three independent biological samples or three independent cell cultures), and a representative set of data was used to prepare the figures.

### Co-immunoprecipitation (Co-IP) Assays

Co-immunoprecipitation (Co-IP) assays were performed as described previously [46]. S2 cell lysates (300 μl, approximately equivalent to 10^6^ cells) or equivalent cell culture media containing recombinant proteins were precleared with Protein G Sepharose (50% slurry, No.17-0618-01, GE Healthcare) prior to Co-IP assays. Cell lysates or cell lysates combined with cell culture media were mixed with anti-Flag M2 or anti-V5 antibody (final concentration of 1 μg/μl), and these mixtures were incubated at 4^°^C for 10 h with gentle rocking. Protein G Sepharose (30 μl of 50% slurry) in lysis buffer was added to the protein-antibody mixture, and the resulting mixture was incubated overnight at 4^°^C with gentle rocking. The Sepharose beads containing immunoprecipitated proteins were collected after centrifugation, washed three times with lysis buffer, resuspended in 50 μl of 1 × SDS sample buffer, boiled at 95^°^C for 5 min, and used for subsequent Western blot analysis using anti-Flag M2 or anti-V5 antibody as the primary antibody as described above [46].

Co-immunoprecipitation (Co-IP) assays were also performed by mixing S2 cell culture medium containing Toll^ecto^ or Toll-7^ecto^ proteins collected 48 h after protein expression with DMEM cell culture medium from VSV-GFP-infected HEK293 cells (containing VSV-GFP virions) as described above, and the interaction of Toll^ecto^ or Toll-7^ecto^ with VSV-GFP was detected by anti-V5 or anti-VSV glycoprotein (anti-VSV-G) antibody [P5D4] (ab50549, Abcam, USA, 1:5000 dilution).

### Western blot analysis

Western blot analysis of transiently transfected S2 cells or stable S2 cell lines (5×10^6^ cells/well) was performed in six-well plates 48 h after the induction of protein expression by copper sulfate (final concentration of 250 μM). The cell culture media (2 ml each) and S2 cells were collected, and the S2 cells were homogenized in 400 μl of lysis buffer (50 mM Tris-HCl, pH 7.4, 150 mM NaCl, 5 mM EDTA, 1% NP-40, and 0.5 mM PMSF) containing protease inhibitor cocktail (P8340, Sigma-Aldrich) following a previously described protocol [46]. The cell homogenates were sonicated briefly and centrifuged, and the supernatants (cell lysates) were collected. The cell culture media (10 μl from a total volume of 2 ml) and cell lysate (10 μl from a total volume of 400 μl, equivalent to ∼5×10^4^ cells) were separated by 8%, 12% or 15% SDS-PAGE, and the proteins were transferred to nitrocellulose membranes (162-0097, Bio-Rad) for Western blot analysis using anti-Flag M2 antibody (F-1804, Sigma-Aldrich, 1:5000 dilution) or anti-V5 antibody (V-8012, Sigma-Aldrich, 1:5000 dilution) as the primary antibody and alkaline phosphatase-conjugated anti-mouse antibody (A4312, Sigma-Aldrich, 1:10,000) as the secondary antibody as described previously [47]. The signal was developed using an Alkaline Phosphatase (AP)-Conjugate Color Development Kit (#170-6432, Bio-Rad).

### Real-time PCR analysis

The total RNA from flies and S2 cells was extracted, and the expression of target genes was determined by real-time PCR as described previously [37]. The flies were anesthetized on a CO_2_ bed, placed in 1.5-ml tubes and homogenized with disposable pestles in 1 ml of TRIzol® Reagent (T9424, Sigma-Aldrich), and the total RNA from flies and S2 cells was extracted according to the manufacturer’s instructions. The RNA pellets were air-dried and resuspended in 50 μl of nuclease-free water, and the concentration of RNA was determined using a Nanodrop UV-Vis spectrophotometer (ND-1000, Thermo).

Total RNA (2 μg from each sample) was treated with RQ1 RNase-free DNase (M6101, Promega) to remove contaminated genomic DNA and then used for the synthesis of cDNAs in 25 μl reactions using Moloney murine leukemia virus (M-MLV) reverse transcriptase (M1701, Promega) and an anchor-oligo(dT)18 primer following the manufacturer’s instructions. The cDNA sample (diluted 1:50) was used as the template for quantitative real-time PCR analysis. The *Drosophila* ribosomal protein 49 (*rp49*) gene was used as an internal standard to normalize the expression of target mRNA. Real-time PCR was performed in 20 μl reactions containing 10 μl of 2×SYBR® GreenER™ qPCR SuperMix Universal (No. 204141, Qiagen), 4 μl of H_2_O, 4 μl of diluted cDNA template, and 1 μl (10 pmol) of each of the forward and reverse primers. The real-time PCR program was 2 min at 50^°^C, 10 min at 95^°^C, 40 cycles of 95^°^C for 15 s and 60^°^C for 1 min, and the dissociation curve analysis. The data from three replicates of each sample were analyzed with a comparative method (2^-ΔΔCT^) using ABI 7500 SDS software (Applied Biosystems). The baseline was automatically set by the software to maintain consistency. The cDNA sample from S2 cells transfected with empty pMT/BiP/V5-His A plasmid or wild-type flies (*w*^*1118*^) was used as the calibrator. The expression level of target genes was calculated by the 2^-ΔΔCT^ method [48], which provides the n-fold difference in relative expression compared with the calibrator. All the data are presented as relative mRNA expression levels, and all the experiments were repeated at least three times.

### Data analysis

Three to four replicates of all the experiments were performed, and the experiments were repeated with three to four independent biological samples. The means from a typical dataset were used for the figures, which were prepared using GraphPad Prism (GraphPad, San Diego, California, USA). The statistical significance of the differences was calculated by one-way ANOVA followed by Tukey’s multiple comparison test using GraphPad Prism with identical letters for a non-significant difference (*p* > 0.05) whereas different letters for a significant difference (*p* < 0.05). The significance of the difference was also determined by an unpaired t-test using GraphPad InStat software with * *p* < 0.05, ** *p* < 0.01, and *** *p* < 0.001.

## Acknowledgements

This work was supported by the National Natural Science Foundation of China (No. 31472019) and University of Missouri – Kansas City Funding for Excellence.

## Author contributions

XY designed the experiments, analyzed data, interpreted results, and participated in manuscript writing; MC performed most experiments, analyzed data, interpreted results, and participated in manuscript writing; CL performed some experiments and analyzed data; ZH and YL helped perform some experiments; XL and YW participated in manuscript writing; YI and MS helped interpret results and participated in manuscript writing.

## Expanded View Figure legends

**Figure EV1 - Amino acid sequences of *Drosophila* pro-Spätzle and cistine knot Spätzle proteins, and multiple sequence alignment of *Drosophila* Spz, Spz-2 and Spz-5.**

A The amino acid sequences of pro-Spz to pro-Spz-6 were obtained from the NCBI website (https://www.ncbi.nlm.nih.gov/) with the indicated accession numbers. The predicted cistine knot Spz domains were underlined. B *Drosophila* cistine knot Spz, Spz-2 and Spz-5 domains (from Fig EV1A above) were aligned by Clustal Omega (https://www.ebi.ac.uk/Tools/msa/clustalo/). Identical residues are indicated by “*”, highly conserved residues are indicated by “:”, and conserved residues are indicated by “.”.

**Figure EV2 - Expression of recombinant Toll, Toll-7 and six Spz proteins in S2 cells**.A-D V5-tagged recombinant Toll, Toll-7, Toll^ecto^ and Toll-7^ecto^ as well as Flag-tagged active cistine knot Spz to Spz-6 proteins were expressed in S2 cells. Proteins in the cell culture media (A and B) and the cell lysates (C and D) were detected by anti-V5 (A and C) or anti-Flag (B and D) monoclonal antibody.

**Figure EV3 - Expression of *gfp* transcript in the flies infected with VSV-GFP.**A, B *w*^*1118*^, *Df(2R)BSC22/Toll-7*^*g1-5*^ and *Tlr*^*632*^*/Tl*^*I-RXA*^ male and female flies were infected with VSV-GFP, and flies at 1, 5 and 10 days post-infection were collected for preparation of total RNAs. Replication of VSV-GFP in the flies was determined by expression of *gfp* transcript in the RNA samples by real-time PCR using *Drosophila* ribosomal protein 49 (*rp49*) gene as an internal standard, and relative expression of *gfp* in *w*^*1118*^ males (A) or females (B) was arbitrarily set as 1. No significant difference in the expression level of *gfp* among *w*^*1118*^, *Df/Toll-7*^*g1-5*^ and *Tlr*^*632*^*/Tl*^*I-RXA*^ males (A) and female (B) was observed at days 1, 5 and 10 post-infection, and no significant difference in *gfp* expression level was observed in *w*^*1118*^, *Df/Toll-7*^*g1-5*^ or *Tlr*^*632*^*/Tl*^*I-RXA*^ flies between days 1, 5 and 10 post-infection.

Data information: In (A, B), graphs show mean ± SEM, n=3.

**Figure EV4 - Expression of *Toll* and *Toll-7* transcripts in *w*^*1118*^ and mutant flies after *E. faecalis*, *P. aeruginosa*, *C. albicans*, and VSV-GFP infection.** A Expression of *Toll* and *Toll-7* transcripts in the un-infected *w*^*1118*^ males and females. Real-time PCRs were performed using *Drosophila* ribosomal protein 49 (*rp49*) gene as an internal standard, and expression of *Toll* in *w*^*1118*^ males was arbitrarily set as 1. B-I Expression of *Toll* and *Toll-7* transcripts in *w*^*1118*^, *Toll-7*^*g1-5*^*/CyO*, *Df(2R)BSC22/Toll-7*^*g1-5*^, *T1*^*I-RXA*^/*TM6B* and *Tlr*^*632*^*/Tl*^*I-RXA*^ mutant males (B-E) and females (F-I) after *E. faecalis*, *P. aeruginosa*, *C. albicans* and VSV-GFP infection. Real-time PCRs were performed using *Drosophila rp49* gene as an internal standard and expression of *Toll* or *Toll-7* mRNA in *w*^*1118*^ flies after infection was arbitrarily set as 1. Data Information: In (A-I), graphs show mean ± SEM, n=3; identical letters are not significant difference (*p* > 0.05), whereas different letters indicate significant difference (*p* < 0.05), one-way ANOVA followed by Tukey’s multiple comparison test; * *p* < 0.05, ** *p* < 0.01 and *** *p* < 0.001, unpaired t-test.

**Figure EV5 - Expression of *drosomycin* and *diptericin* transcripts in *w*^*1118*^ and mutant flies after *E. faecalis*, *P. aeruginosa*, *C. albicans*, and VSV-GFP infection.**

A-H Expression of *drosomycin* (*Drs*) and *diptericin* (*Dpt*) transcripts in *w*^*1118*^, *Toll-7*^*g1-5*^*/CyO*, *Df(2R)BSC22/Toll-7*^*g1-5*^, *T1*^*I-RXA*^/*TM6B* and *Tlr*^*632*^*/Tl*^*I-RXA*^ mutant males (A-D) and females (E-H) after *E. faecalis*, *P. aeruginosa*, *C. albicans* and VSV-GFP infection. Real-time PCRs were performed using *Drosophila rp49* gene as an internal standard and expression of *Drs* or *Dpt* in *w*^*1118*^ flies after infection was arbitrarily set as 1.

Data information: In (A-H), graphs show mean ± SEM, n=3; identical letters are not significant difference (*p* > 0.05), whereas different letters indicate significant difference (*p* < 0.05), one-way ANOVA followed by Tukey’s multiple comparison test.

